# Injury-induced transcription in the planarian outer epithelium is critical for tissue regeneration

**DOI:** 10.1101/2024.10.01.616145

**Authors:** Pallob Barai, Mariya S. Kibtiya, Nathan G. Maggard, Shishir Biswas, Elizabeth M. Duncan

## Abstract

Planarian flatworms have an extraordinary regenerative capacity; even a small, asymmetrical fragment of amputated planarian tissue can recreate an entirely new animal. The planarian field has made significant progress in identifying specific genes and cell types required for this complex process, but substantially less is known about the molecular mechanisms that convert a significant physical injury (e.g., head amputation) into the expression of specific genes in particular cell types. One tissue in which this question is particularly relevant is the planarian epidermis, a single-layer, mucociliary epithelium with similarities to the epithelia of mammalian airways. This epithelium plays an essential, early role in planarian regeneration as the cells surrounding the wound site must quickly stretch and extend to cover the wound area after injury. We hypothesized that these injury-induced morphological changes activate the transcription of genes with essential functions in the regenerative process. To best detect these transcriptional changes, we developed a rapid method for isolating the planarian outer epithelium and prepared ribodepleted RNA-sequencing libraries from samples isolated at multiple time points after tissue amputation. One gene we both identified using these methods and found to be functionally important for regeneration is a putative planarian homolog of vertebrate *Shoc2*. SHOC2 is an essential scaffolding protein that mediates specific, context-dependent activation of the ERK1/2 signaling pathway (J ang andG alperin2016). Notably, the ERK1/2 pathway is known to be activated after injury and required for regeneration in multiple species (M anuel *et al*. 2006; O wlarn*et al*. 2017; T omasso*et al*. 2023). These findings suggest that these epithelial datasets have the potential to uncover many functionally relevant and possibly highly conserved genes that play fundamental roles in animal regeneration.

## INTRODUCTION

The outer epithelium of planarian flatworms acts as both a mucociliary epithelium, sharing many features with human airway epithelia (R ompolas*et al*. 2009), and as a simple epidermis that provides a barrier to the external environment. Planarians rely on both the mucous-secreting and multiciliated cells of this epithelium for their motility, as the beating motion of the dense cilia propel them along a mucous-paved path. This outer epithelium is also critical for one of the first essential steps of planarian regeneration: rapid wound closure. Immediately after a major injury occurs, the planarian body wall muscle contracts, reducing the surface area of the wound, and the epithelial cells around the wound site flatten and extend to cover the exposed tissue (C handebois1979; H ori1989). Together, these changes are necessary for regeneration to proceed, as inhibiting muscle contraction prevents subsequent blastema formation (S chÜrmann1998). However, it is unclear whether the morphological changes that occur in the planarian epithelium are specifically critical to the regenerative process.

Studies in mammalian cells and other models have shown that injury-induced mechanical stress can activate relevant changes in gene expression (C opland andP ost2007; H ellman*et al*. 2010; G udipaty*et al*. 2017; H irashima*et al*. 2023). We therefore hypothesized that the physical changes induced in the planarian epithelium after injury are likely to initiate changes in gene expression that are critical for the subsequent steps of tissue regeneration. To test this hypothesis, we first developed a robust method for rapidly and effectively removing the outer epithelium (of *Schmidtea mediterranea, or S*.*med*, specifically), as a previously published method involved soaking planarians in high salt for a significant amount of time (W urtzel *et al*. 2015). Such procedures are not optimal for measuring gene expression changes at early time points after injury, nor are they compatible with downstream genomic assays (e.g., ChIP-seq, ATAC-seq) as the salt solution will likely disrupt the chromatin state. After establishing that our method worked well for removing the outer epithelium and extracting high quality RNA from the separated tissue, we then used it to identify transcriptional changes occurring in the outer epithelium at early stages of regeneration. Further, we used a recently optimized method for generating RNA-seq libraries from ribodepleted RNA samples (B arai *et al*. 2024), making our datasets useful for characterizing the expression of both coding and non-coding genes.

In our initial analysis of these datasets, we detected activation of both genes that were previously reported to be induced in the epithelium after injury e.g., *hadrian* (W *enemoser* *et al. 2012*; *W *urtzel* *et al. 2015*), atp1a1* (B enham-P yle *et al*. 2021), and genes that have not yet been characterized. As we screened the latter genes for potential function in regeneration, we uncovered one of particular interest: *Smed-shoc2b*, a putative homolog of vertebrate Shoc2. This gene encodes a scaffolding protein that is essential for ERK activation in the MAPK/ERK signaling pathway (J ang and G alperin 2016; J eoung *et al*. 2016), a pathway that is known to be activated and required for wound healing and regeneration in several different species (M anuel *et al*. 2006; O wlarn *et al*. 2017). RNAi of *Smed-shoc2b* revealed that this putative homolog plays a functional role in both planarian regeneration and homeostasis, although it is not yet clear if they are mediated through the same cell type and/or mechanism. Nonetheless, this finding has raised exciting questions about the possibility of a conserved role for Shoc2 in regeneration, particularly as a recent study found that ERK activation is sustained longer after in the regenerative spiny mouse versus non-regenerative house mouse (T omasso *et al*. 2023). It also supports our assertion that the methodology and data we report here have the potential to uncover many genes that are both essential and conserved in animal regeneration.

## RESULTS

### A new method for rapidly isolating the planarian outer epithelium

To identify genes that undergo significant changes in expression in the outer planarian epithelium after injury, we first developed and optimized a method for quickly removing the epithelium (**Figure 1a**). After studying the literature and trying various methods, we found that a simple solution of 0.1% SDS (in Montjuïc water, see Methods) worked effectively to quickly loosen the outer planarian epithelium from the underlying tissue (**Figure 1b**) allowing it to be gently and cleanly peeled away from the rest of the worm with a syringe needle (**Figure 1c**). To verify that we successfully isolated the outer epithelium cleanly and in a manner compatible with gene expression studies, we extracted its total RNA, used it to generate cDNA, and then performed qPCR to measure the expression of the following lineage-specific marker genes: *piwi-1* (stem cells), *prog-1* (early progenitors), *agat-1* (late progenitors), and *rootletin* (ciliated cells in the outer epithelium). When we compared the expression of these marker genes in isolated epithelial tissue (epi) and whole planarian worms (WW), we found that only *rootletin* was enriched in the outer epithelium (**Figure 1d**) and detected little to no expression of the other markers in this tissue, as expected.

**Figure 1.**
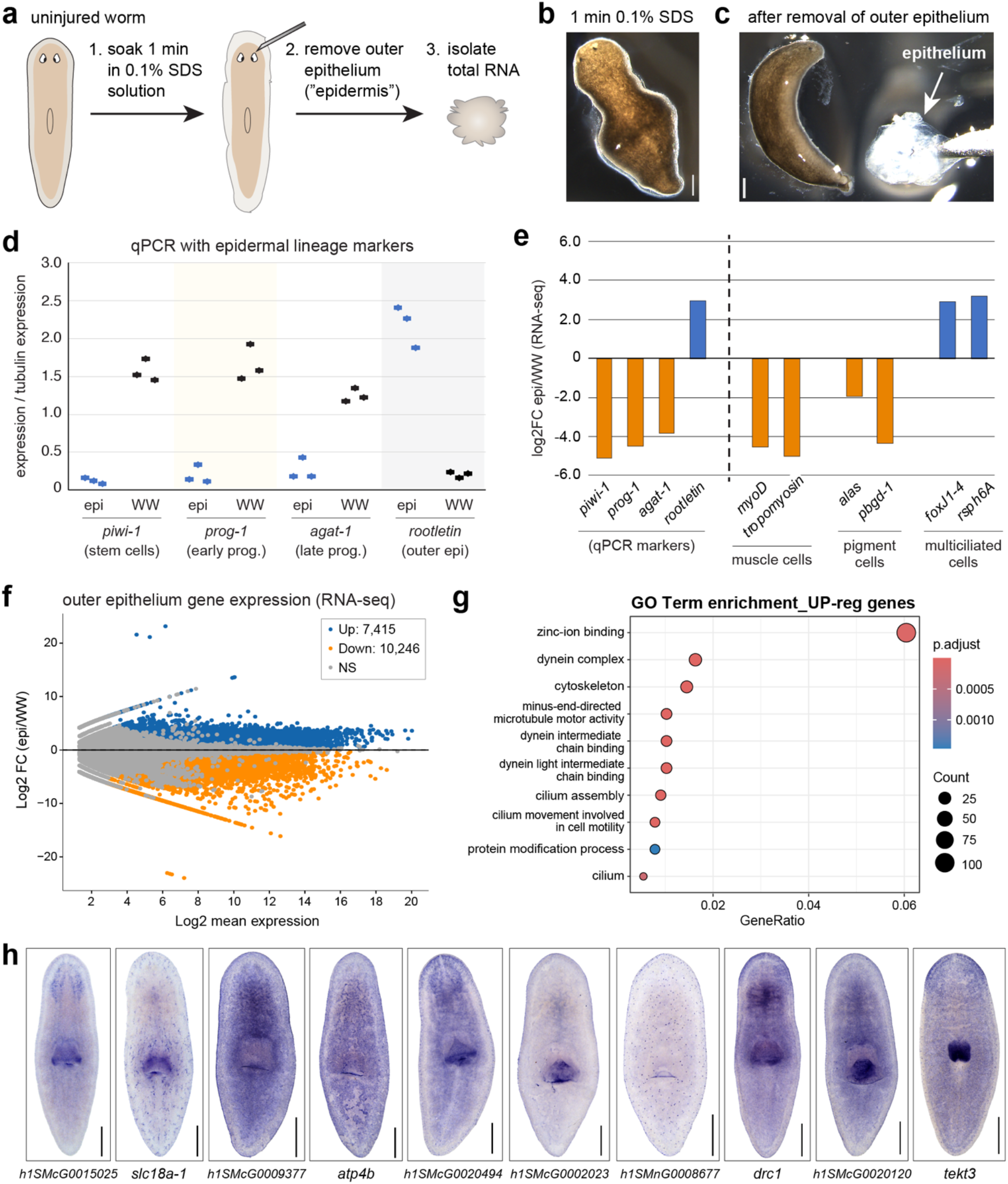
A new method for removing the planarian outer epithelium is rapid and precise. a) Cartoon illustrating method for the removal of the planarian outer epithelium (a.k.a. “epidermis”) using a 0.1% SDS solution and surgical needle. b) image of live planarian worm after 1 minute incubation in 0.1% SDS solution at room temperature. c) image of live planarian worm after removal of outer epithelium; white arrow indicates epithelium after removal; rest of worm (left) remains largely intact. d) Quantitative PCR (qPCR) assessing the relative expression of markers genes for the epidermal lineage, including both those that are markers for stem cells and epithelial progenitor cells (*piwi-1, prog-1, agat-1*) and one that is a marker of the multiciliated cells of the outer epidermis (*rootletin*). Data is shown for three independent samples of isolated epidermal tissue (epi) and whole planarian worms (WW). e) Plot showing the relative expression of representative transcripts in RNA-seq data generated from epidermal tissue (epi) and whole planarian worms (WW). f) MA plot of RNA-sequencing data from isolated outer epithelium samples (epi) compared with whole planarian worms (WW). Blue dots represent genes significantly up-regulated in the epithelium (DEseq2) with a log2FC > 1.0 and pAdj <0.01. Orange dots represent genes with relatively less expression in the epithelium compared to whole worm tissue (DEseq2), log2FC < -1.0 and pAdj <0.01. g) GO Term enrichment analysis for those transcripts identified as epi enriched in f. h) Whole Mount In Situ Hybridization (WISH) on uninjured, wild-type planarians using riboprobes for transcripts identified as epi enriched in f. Scale bars in b, c, and h = 500 µm.

After establishing that this new method worked well for assessing gene expression in the planarian outer epithelium, we used the total RNA to generate comprehensive RNA-seq libraries using a customized ribodepletion method (B arai *et al*. 2024). After analyzing these data using the newest version of the *S*.*mediterranea* genome (I vankovic *et al*. 2024), we first wanted to confirm that this RNA-seq data showed the same pattern of marker gene expression as detected using qPCR. Indeed, we found that genes associated with multiciliated cells (*rootletin, foxJ1-4, rsph6A*) were enriched in the RNA-seq libraries made from the outer epithelium (epi) whereas gene markers of stem cells (*piwi-1*), epidermal progenitors (*prog-1, agat-1*), muscle cells (*myoD, tropomyosin*), and pigment cells (*alas, pbgd-1*) were significantly better represented in the libraries made from whole worm RNA (WW; **Figure 1e**).

We next wanted to assess the range of transcript detection and Gene Ontology (GO) terms associated with those genes found to be enriched in the outer epithelium. We found that those genes significantly enriched in the outer epithelium (blue) span a broad range of expression levels (**Figure 1f**). We then performed GO Term enrichment analysis on those genes significantly enriched in the outer epithelium (up-regulated with a pAdj <0.01) and found that 7/10 enriched GO Terms were associated with cilia (**Figure 1g**). Upon closer inspection, we realized that the genes in the category “protein modification process” are also likely related to cilia as they were all enzymes that modify tubulin (**Supplemental Table S1**). Lastly, we wanted to visualize the overall expression patterns of representative epi-enriched genes. We performed Whole Mount In Situ Hybridization (WISH) on uninjured planarians for transcripts with a range of expression levels (**Figure 1f, h**). Several of these genes showed strong, broad expression in the outer epithelium (e.g., *atp4b*), some showed a pattern suggesting expression across multiciliated cell types (e.g., *tekt3*), and others suggested restricted expression in a subset of epidermal cells (e.g., unknown transcript *h1SMnG0008677*). Together, these findings strongly support the conclusion that our new method for the rapid removal of the planarian epithelium is highly useful for identify genes expressed in this tissue, including lowly expressed genes.

### Injury induces significant transcriptional changes in the outer epithelium

After establishing that our new method can rapidly remove the planarian outer epithelium and generate comprehensive RNA-seq libraries, we wanted to use it to characterize the gene expression changes that occur in this tissue after injury. Because we wanted to assess transcriptional changes, not changes in tissue composition, we decided to isolate epithelial tissue from trunk fragments at early time points after amputation (10 minutes, 1 hour and 3 hours) and compare each to 24 hours post-amputation (**Figure 2a**). For each time point we pooled the epithelia of 8 trunk fragments and generated three independent replicates (24 worms total per time point). We then extracted total RNA from these samples and generated ribodepleted RNA-seq libraries to allow for the robust detection of both coding and putative non-coding transcripts. After analyzing this sequencing data using the most recent *S*.*mediterranea* genome (I vankovic *et al*. 2024), we identified over 1000 genes with significant (pAdj < 0.01) differential expression at one or more time points (**Figure 2b; Supplemental Table S2**).

**Figure 2.**
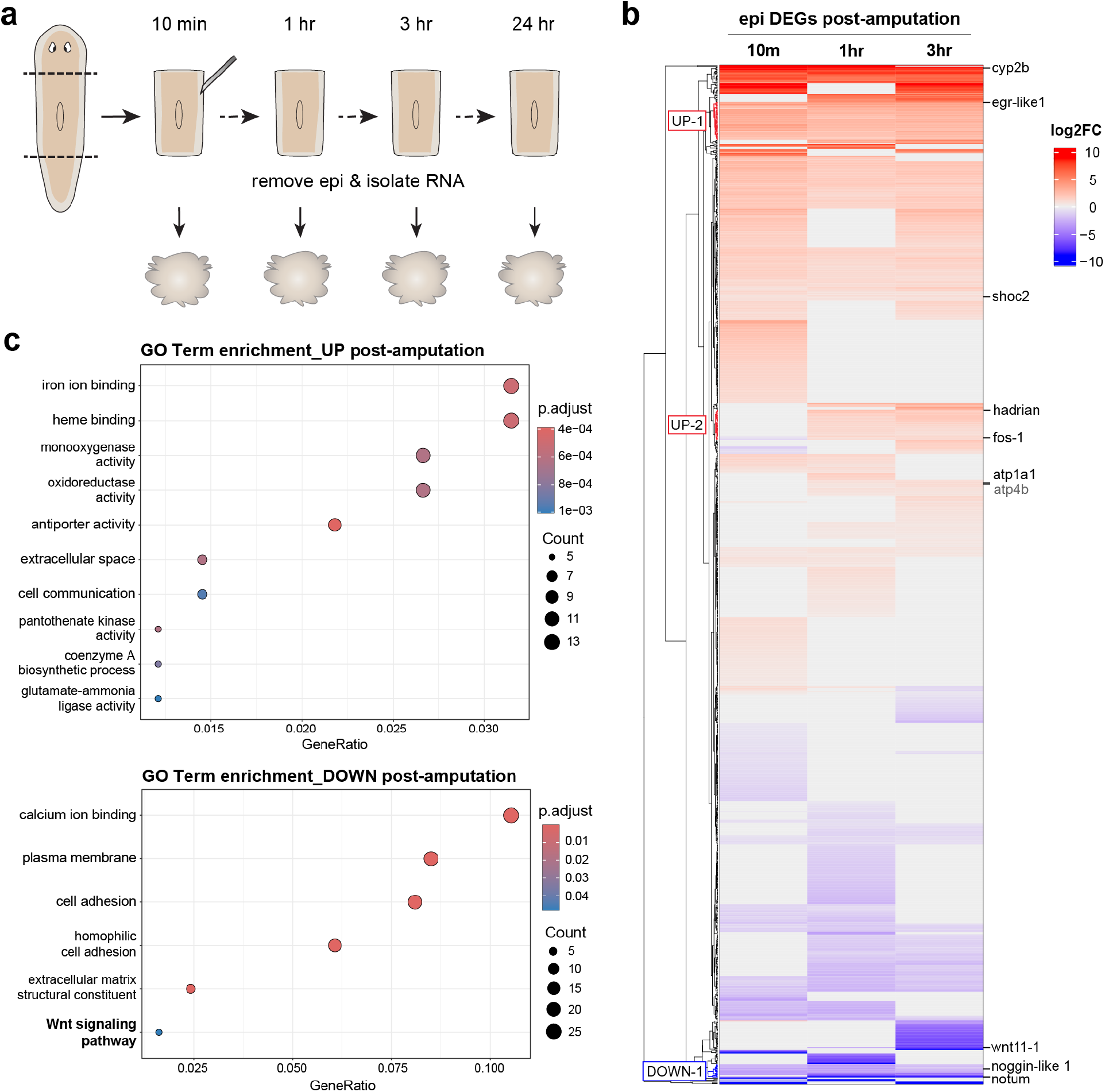
RNA-seq reveals transcripts that are activated in the outer epithelium after major injury. a) Cartoon illustrating the experimental schema, designed to identify significant changes in transcription rather than tissue composition. b) Heat map of differentially expressed genes (DEGs) at the indicated time points after tissue amputation. Each time point (10m, 1hr, 3hr) is compared to the 24hr samples. Genes were included if they had a significant change in expression (pAdj <0.01) for at least one time point. Select subclusters are highlighted for further analysis in Figure 3. C) GO Term enrichment analysis for genes with significant up-regulation after injury (i.e., top part of heatmap) and those with significant down-regulation after injury (i.e., bottom part of heatmap) as compared to the 24hr samples.

Reassuringly, these genes included several genes that were previously reported to be upregulated after injury, including *egr-like1, hadrian, fos-1*, and *atp1a1* (S andmann *et al*. 2011; W enemoser *et al*. 2012; W urtzel *et al*. 2015; B enham-P yle *et al*. 2021).

To develop a better understanding of what types of genes and processes are up or down-regulated in the outer epithelium after injury, we performed Gene Ontology (GO) Term enrichment analyses on both the upregulated and down-regulated genes (**Figure 2c and Supplemental Table S2**). Several of the GO Terms enriched in the up-regulated gene set are associated with the activation of cytochrome P450 genes (**Figure 2b, c**). Others, not unexpectedly, are related to functions related to intercellular communication. Notably, the list of significantly “down-regulated” genes and GO terms includes *notum* and other Wnt signaling pathway components (**Figure 2b, c**). Although we reasoned that they were likely calculated as “down-regulated” because their expression peaks around 24 hours, the time point we used as our denominator, we were not expecting to detect them in epithelial cells as at least some of these genes are reported to be expressed in muscle cells (W itchley *et al*. 2013; S cimone *et al*. 2017; G ittin and P etersen 2022). However, many of those papers use double Fluorescent In Situ Hybridization (FISH) with muscle marker genes to determine this expression, which may not detect expression (especially weaker expression) in other cells types. Moreover, the expression of *notum*, for example, is less clearly restricted to muscle cells in single cell RNA-seq ATLAS data (F incher *et al*. 2018; P lass *et al*. 2018). It is also possible that because the wound epithelium at the injury site is thinner and not secured to a basement membrane, neighboring cells are removed along with it after 0.1% SDS treatment. As transcriptional changes in these neighboring cells are also of interest, their presence in the dataset is additive but something that must be considered and addressed in follow-up experiments.

We then wanted to examine the gene expression changes in this post-amputation time course more granularly. For example, we wanted to substantiate our hypothesis that many “down-regulated” genes in fact have increased expression at or around 24 hours. We chose three gene clusters in the DEG heatmap (“UP-1”, “UP-2, and “DOWN-1” in **Figure 2b**) to examine using Transcripts per kilobase Million (TPM). The genes in all three of these clusters all generally correlate with their log2FC trends (**(Figure 3a-c**). The “UP-1” cluster, which includes a gene that was previously reported to be injury-induced (egr-like1 (S andmann *et al*. 2011)), is a group of genes that are activated immediately and stay upregulated through at least 3hr (**Figure 3a**). The “UP-2” cluster, which includes two genes previously reported to be injury induced (fos-1 and hadrian (W enemoser *et al*. 2012; W urtzel *et al*. 2015), includes genes that are activated slightly later (1hr) but then stay upregulated through 3hr (**Figure 3b**). Analysis of the “DOWN-1” cluster, which includes both *notum* (*P* *etersen and* *R* *eddien* *2011*) and *noggin-like-1* (M olina *et al*. 2011), indeed shows that most of these genes have very little expression at the early time points (10m, 1hr, 3hrs) but show a significant increase in expression at 24 hrs (**Figure 3c**). The genes in this “DOWN-1” cluster all have very low expression in the intact epithelium as well (**Supplemental Table S2**). These transcriptional patterns are also observed by WISH for some genes with these TPM dynamics (**Figure 3d-f**).

**Figure 3.**
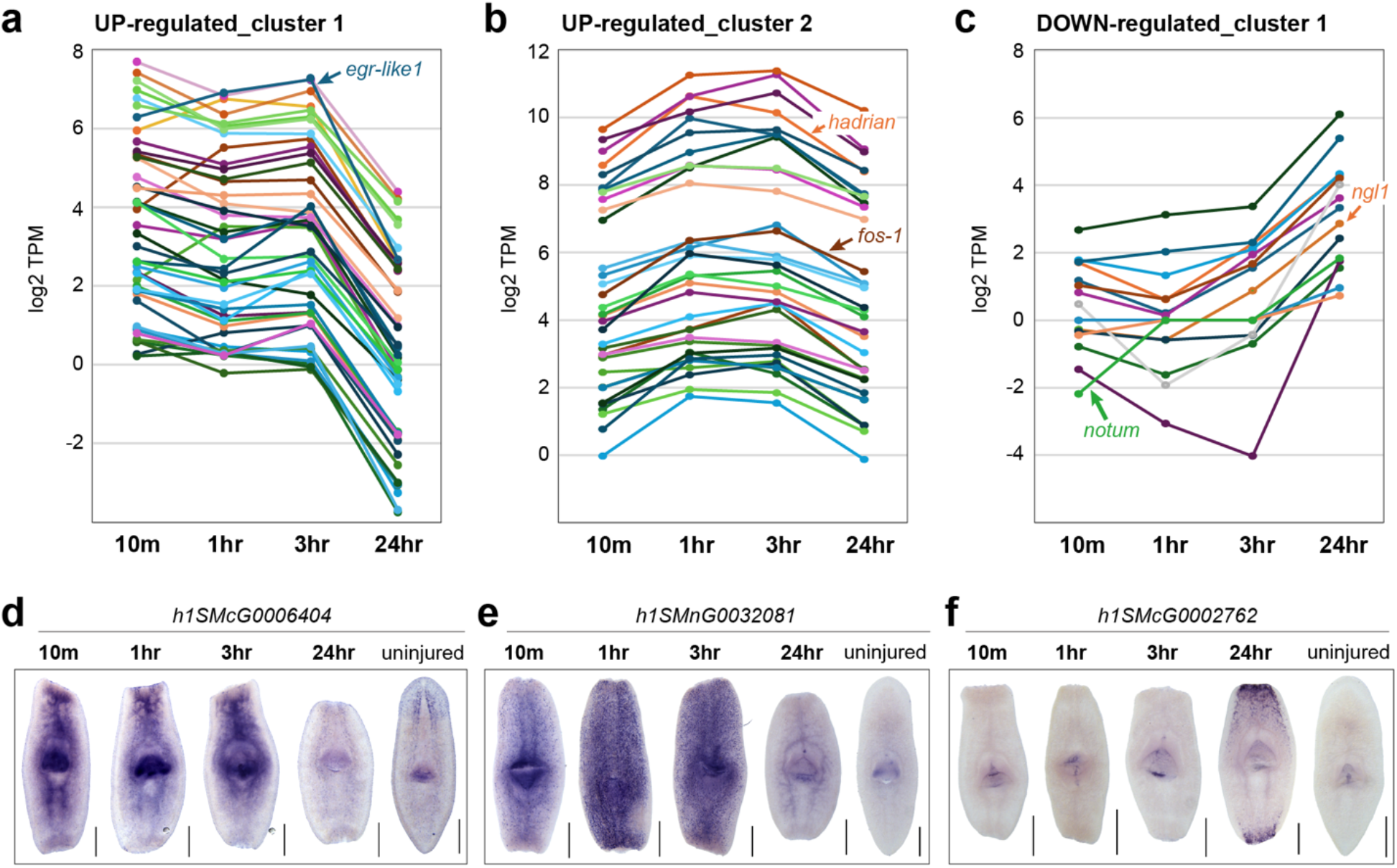
Injury-induced transcriptional changes cluster according to their dynamics. a) Plot showing the expression changes (in TPM, or Transcripts per kilobase Million) for genes labeled in Figure 2b as “UP 1” b) “UP 2”, and c) “DOWN 1”. d) Whole Mount In Situ Hybridization (WISH) was performed on regenerating trunk fragments for genes reflecting the dynamics shown in a-c. Scale bars = 500 µm.

### Genes activated in the outer epithelium are important for regeneration

After identifying many genes with significant differential expression, we wanted to determine if any have important functional roles in regeneration. We use RNA interference (RNAi) to knockdown each planarian gene of interest, split the treated worms into two groups (regeneration or homeostasis), and amputated those animals in the “regeneration” group with transverse cuts above and below the pharynx (**Figure 4a**). As of now, we have screened approximately 100 genes (identified in **Figure 2**) and identified several with reproducible phenotypes (**Supplemental Table S3**). One gene that revealed an RNAi phenotype was shoc2b, a potential homolog of vertebrate Shoc2 (**Figure 4b-d**). This gene drew our attention because its putative homolog encodes a scaffolding protein required for MAPK activation of ERK (J ang and G alperin 2016; J eoung *et al*. 2016), and ERK activation is required for regeneration in multiple species (M anuel *et al*. 2006; O wlarn *et al*. 2017; T omasso *et al*. 2023). RNAi of *S*.*med-shoc2* caused regenerating fragments to develop significantly smaller blastemas (**Figure 4b,c**) with significantly delayed or no new eyespot formation at 8 days post amputation (**Figure 4b**). In addition, we observed that both regenerating head fragments (**Figure 4b**) and uninjured worms (**Figure 4d**) gradually lost their eyespots over time but regained them if we stopped RNAi treatment. Together these data suggest that the regulation of *shoc2b* transcription may play an important role in the dynamic regulation of ERK signaling (**Figure 5**).

**Figure 4.**
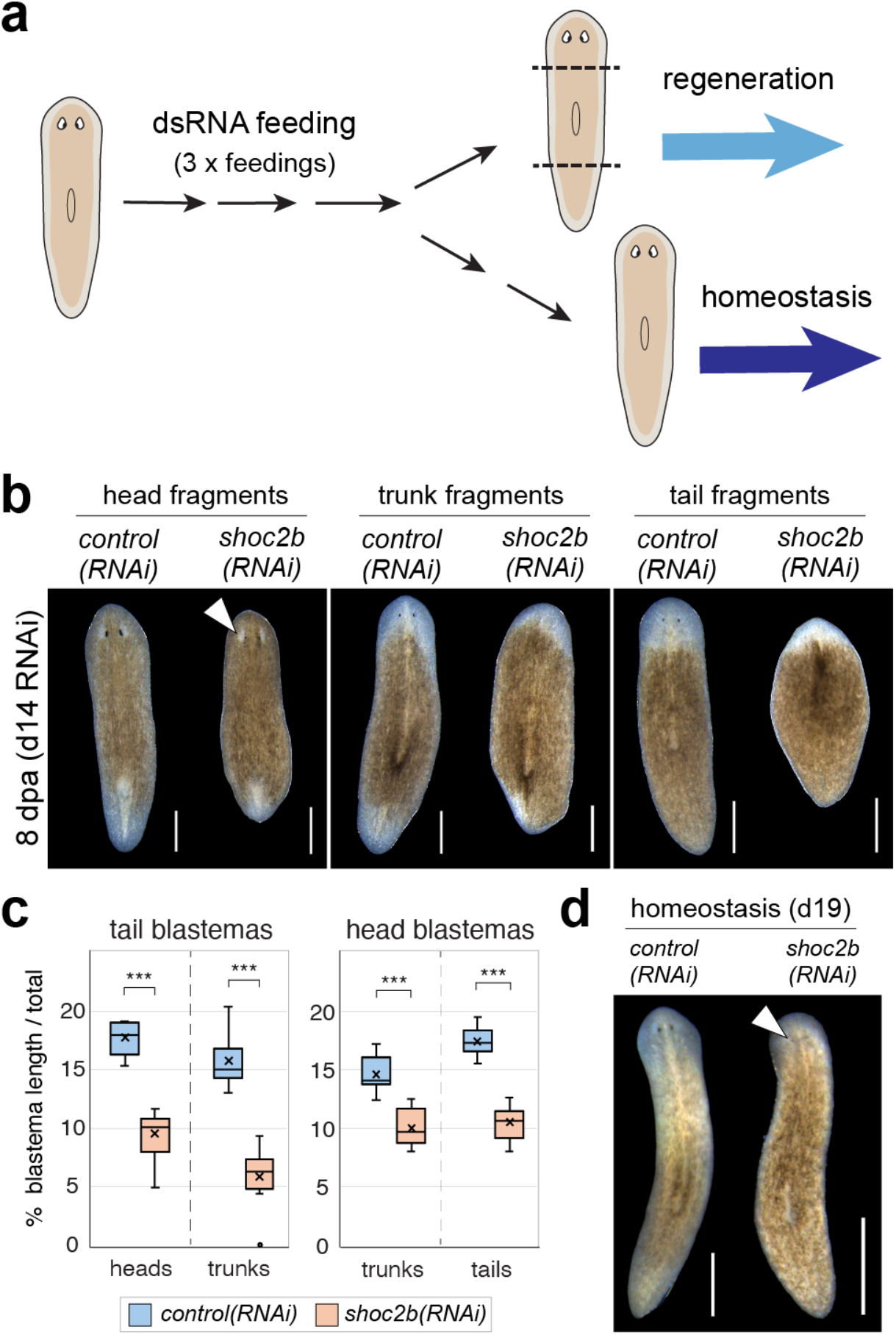
Expression of shoc2b is important for normal regeneration and maintenance of planarian eyespots. a) Cartoon illustrating the experimental schema, designed to identify gene function in planarian tissue regeneration and/or homeostasis. b) Live images comparing the effects of *shoc2b(RNAi)* to *control(RNAi)* worms at 8 days post amputation (8dpa). c) Plots summarizing the relative size of regenerating tail blastemas (left) and head blastemas (right) for all worms included in the experiment in b. d) Live images comparing *control(RNAi)* and *shoc2b(RNAi)* in uninjured worms (homeostasis). White arrowheads point to the disappearing eyespots *shoc2b(RNAi)* animals. Scale bars in b = 200 µm, in d = 500 µm.

**Figure 5.**
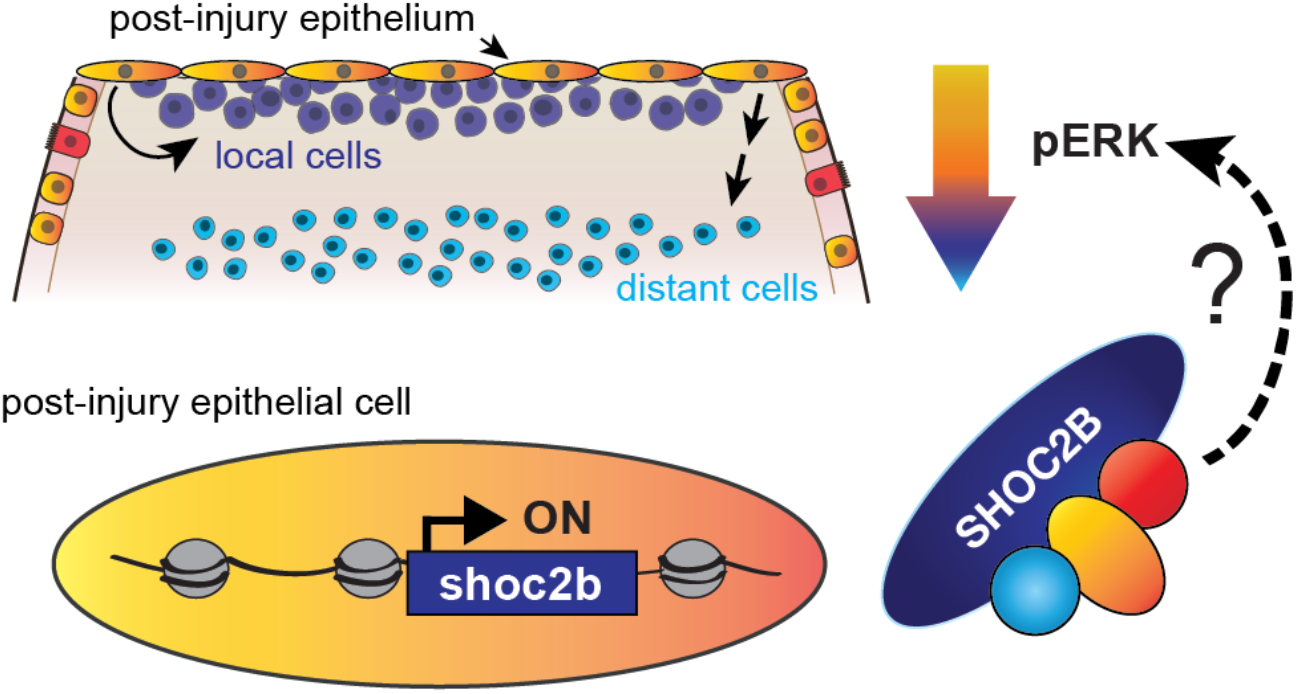
Increased transcription of shoc2b may be required for the maintenance and propagation of ERK activation during tissue regeneration. In this model, the morphological changes induced in the planarian outer epithelium after acute injury (e.g. decapitation) produce signals that affect both local cells and, indirectly, those further from the injury site. One important change is the upregulation of shoc2b, a potential homolog of vertebrate SHOC2, which is a scaffolding protein in the MAP kinase signaling pathway and required for ERK activation. Sustained activation of ERK (i.e., phosphorylation of ERK) was recently shown to be important for tissue regeneration in mammals (T omasso*et al*. 2023). This model proposes that increased expression of SHOC2 helps mediate sustained ERK activation.

## DISCUSSION

Tissue regeneration, in planarians and other regenerative animals, is a remarkable and complex phenomenon. Although significant progress has been made in identifying many genes that are essential to this process, many questions remain about the molecular mechanisms through which they operate, and how these mechanisms are coordinated in across cell types and time points. In the planarian field, one challenge in making these connections has been our inability to isolate specific cell and/or tissue types of interest. Recent advances in single cell approaches (both RNA-seq and ATAC-seq) will certainly prove useful in addressing these questions, but these methods still present many barriers, including their high cost. Here we present a simple, quick, and inexpensive method for removing the planarian outer epithelium, a tissue with both important and highly conserved functions. Not only is this method useful for measuring changes in gene expression but, unlike other methods for isolating this tissue (W urtzel *et al*. 2015), it also uses a solution (0.1% SDS) that is compatible with genomic assays such as ChIP-seq.

Epithelial tissue isolated with this method can also be used for protein isolation and analysis, particularly for applications like immunoblotting.

We sought to develop a method for isolating the outer epithelium because it stands out as a tissue with a clear role in the early stages of regeneration and yet relatively little knowledge about the molecules and mechanisms mediating this role. Unsurprisingly, we are not the first group to identify the importance of this tissue and its role in regeneration. Nonetheless, there have been relatively few genes identified as both activated in the epithelium after injury and functionally important in planarian regeneration(W enemoser*et al*. 2012; W urtzel *et al*. 2015; S cimone*et al*. 2022). Here we report multiple RNA-seq datasets that have the potential to uncover many novel and/or uncharacterized transcripts with important roles in tissue regeneration. We also show a powerful example of a gene that fits this model, *Smed-shoc2b*. Recent work in planarians and other organisms have collectively focused on the functional importance of ERK activation, its propagation (F an*et al*. 2023), and its timing (T omasso*et al*. 2023) in tissue regeneration. However, the details of how ERK is activated in response to injury, the tissues in which this occurs, and the mechanisms mediating its dynamics are unclear. We propose a model in which the injury-induced upregulation of *shoc2* mediates the maintenance, and possibly the propagation, of ERK activation and signaling (**Figure 5**). We are currently testing this hypothesis as part of the many future directions of the work presented here.

## METHODS

### Planarian maintenance and care

Asexual strain of Schmidtea mediterranea (clonal line CIW4, NCBI Taxon: 79327) (S anchez A lvarado *et al*. 2002) were maintained in 1X Montjuic water (1.0 mmol/l calcium chloride, 1.6mmol/l sodium chloride, 1.0mmol/l magnesium sulfate, 0.1mmol/l magnesium chlorid, 0.1 mmol/l potassium chloride and 1.2 mmol/l sodium bi carbonate in Milli-Q water, pH-6.9-8.1) (C ebria andN ewmark2005). Animals were fed beef liver paste and worms were kept starved for a week before using them in any experiments.

### Planarian epidermis isolation and RNA extraction

Before isolating the epidermis, 6-8mm worms were kept starved for a week. Intact or amputated worms (10 minutes post amputation, 1 hour post amputation, 3 hpa and 24 hpa) were soaked in 0.1% SDS for 1 minute on a glass plate. After soaking, the epidermis of the worms was gently peeled off using a 21G syringe needle while visualizing under a stereo microscope. The tissue was transferred in 0.5ml of TRIzol reagent kept on ice. For each sample, the skin of 8-10 worms or fragments were pooled into one tube. RNA extraction for samples in TRIzol were performed using Zymo Research RNA clean and concentrator-5 kit according to manufactures protocols with minor variation. For extracting the RNA, we used size selection before further RNA-seq library preparation as previously described (K im *et al*. 2019b).

### RNA extraction and ribodepletion with RNaseH

Total RNA was extracted from whole planarian worms or regenerating tissue using TRIzol reagent according to the manufacturer protocol. After extracting RNA, it was further cleaned, concentrated, and size-selected for RNAs > 200nt using the Zymo Research RNA Clean & Concentrator-5 kit (Catalogue No. R1013). An RNase H mediated rRNA depletion protocol was adapted from methods developed for bacteria (C ulviner *et al*. 2020) and human cells (B aldwin *et al*. 2021). Briefly 500 ng of size-selected total RNA was mixed with 1000ng of a species-specific rRNA probe pool (**Supplemental Table S4**) and hybridization buffer (final 50mM Tris HCl pH 7.5, 100mM NaCl, 20mM EDTA) in a final volume of 15µL. Probe-RNA mixtures were heated in a heat block at 95°C for 2 mins and then slowly cooled (-0.1°C/sec) to 65°C. The mixtures were then kept at 65°C additional 5 minutes while the RNaseH mixture was prepared: for each RNA sample, 3μL Lucigen Hybridase™ Thermostable RNase H enzyme (Catalogue No. H39500), 0.5μL 1M Tris-HCl pH 7.5, 0.2μL 5M NaCl, 0.4μL 1M MgCl2, and 0.9μL nuclease-free water (total 5μL) was mixed and preheated to 65°C. The 5μl preheated Hybridase RNase H reaction was then added to the sample mix and incubated at 65°C for 2.5 minutes. In-column DNase I treatment was then performed to degrade excess probes and genomic DNA; 30µL of prepared DNase I reaction mix (3μL TURBO™ DNase [ThermoFisher AM2239] + 5μL10X TURBO™ DNase Buffer + 22μL nuclease-free water) was added to the samples immediately and incubated at 37°C for 30min. The ribodepleted RNA was then cleaned using the Zymo Research RNA clean and concentrator-5 kit (Catalogue No. R1013) and RNA-seq libraries were prepared using the KAPA RNA hyper kit (Catalogue No. 08098107702) along with KAPA unique dual index adapters (Catalogue No. 8861919702) according to the manufacturer protocol.

### RNA sequencing, alignment to the S3h1 genome, and identification of putative ncRNA loci

RNA-seq libraries were shipped to the DNA Services Lab in the Roy J. Carver Biotechnology Center at the University of Illinois Urbana Champaign, checked for quality, and sequenced using either paired-end sequencing technology (intact epithelium and whole worm samples) or single-end technology (epithelial time course) on an SP flowcell of the Illumina NovaSeq 6000. The resulting sequences were then aligned to the S3h1 genome (I vankovic 2023) using HiSat2 (K im *et al*. 2019a). Transcripts per Million mapped reads (TPMs) for each gene model were calculated using Stringtie (P ertea *et al*. 2015b). Differential gene expression analysis was then performed using DESeq2 (version 1.42.0) (L ove *et al*. 2014; P ertea *et al*. 2015a). Further quantitative and statistical analysis was performed using R (version 4.3.1) and figures were created using ggplot2 (W ickham 2009).

### cDNA synthesis for Gene cloning and qPCR

After extracting RNA from whole worms and epidermis, iScripts reverse transcriptase supermix (Biorad Cat #1708840) was used to prepare cDNA. qPCR was performed using BioRad iTaq Universal SYBRGreen mix (Cat #1725120). For cloning the candidate genes, gene specific primers with overhanging nucleotides (forward-CATTACCATCCCG and reverse-CCAATTCTACCCG) homologous to pPR-T4P vector was used. pPR-T4P vector was linearized by treating with SmaI (NEB Cat # R0141L). After treating both the PCR product and linearized vector with T4 polymerase (Novagen Cat #70099), vector and insert were mixed and incubated at room temperature for 60min. Reactions were transformed into E. coli DH5α and positive colonies were cultured for miniprepeps, which were prepared using ZymoPURE Plasmid Miniprep Kit (Zymo Research Cat #D4211). All primers used in this study are listed in **Supplementary Table S5**.

### NBT/BCIP whole mount in situ hybridization

Nitroblue tetrazolium/5-bromo-4-chloro-3-indolyl phosphate (NBT/BCIP) colorimetric whole mount in situ hybridizations were adapted from previously described (G uerrero-H ernÁndez *et al*. 2021) with minor modification. Briefly, 3-5 mm worms were starved 7-10 days prior starting the in situ. Animals were killed in 0.5% HNO3 and fixed with 5% formic acid for 45 mintues. Following the bleaching with 1% formamide and 5% H2O2 solutions for 2 hours, worms were treated with proteinase K (2ug/ml). Animals were hybridized overnight with riboprobes and then samples were washed 2X with wash buffer, 1:1 wash buffer-2X SSC, 3X with 2X SSC, 0.2X SSC and 1X MABT buffer. Subsequently, 0.5% Western Blocking Reagent (Roche Cat # 11921673001) with 5% inactivated horse serum in 1X MABT was used as blocking solutions. Anti-DIG antibody in blocking solutions (1:2500) was used as antibody solutions and samples were incubated in antibody solutions for overnight at 4C. Post antibody washes and colorimetric development was performed as described (K ing and N ewmark 2013).

### Double stranded RNA synthesis and RNAi gene knockdown experiments

In vitro double stranded RNA (dsRNA) synthesis was conducted as previously described (R ouhana *et al*. 2013). Gene knockdown was performed by feeding dsRNA corresponding to the target gene mixed with liver paste. For RNAi experiments, animals were starved 7 days prior to start the dsRNA feeding. Animals were kept in dark for at least 1 hour before each feeding. For each experiment, dsRNA was mixed with homogenized beef liver (1:3) and fed for 3 times in 8 days period. On last day of feeding, animals were cut into head, tail and trunk fragment after 6 hours post feeding and each fragment was allowed to regenerate for 15 days. In this period worm fragments were monitored for every two days for regeneration defects.

### Image processing and quantification

All the live images of live animals and *in situ* hybridizations reported in this study were taken using Leica stereoscope model M205 FA. Images were captured using Leica image processing software LAS X version 3.7.4. We quantified the whole planarian body and blastema size using FIJI (S chindelin *et al*. 2012) and assessed the significance of any differences in blastema size using the Student’s t-test.

